# The correspondence between EMG and EEG measures of changes in cortical excitability following transcranial magnetic stimulation

**DOI:** 10.1101/765875

**Authors:** Mana Biabani, Alex Fornito, James P. Coxon, Ben D. Fulcher, Nigel C. Rogasch

## Abstract

Transcranial magnetic stimulation (TMS) is a powerful tool to investigate cortical circuits. Changes in cortical excitability following TMS are typically assessed by measuring changes in either conditioned motor-evoked potentials (MEPs) following paired-pulse TMS over motor cortex or evoked potentials measured with electroencephalography following single-pulse TMS (TEPs). However, it is unclear whether these two measures of cortical excitability index the same cortical response. Twenty-four healthy participants received local and interhemispheric paired-pulse TMS over motor cortex with eight inter-pulse intervals, sub- and suprathreshold conditioning intensities, and two different pulse waveforms, while MEPs were recorded from a hand muscle. TEPs were also recorded in response to single-pulse TMS using the conditioning pulse alone. The relationships between TEPs and conditioned-MEPs were evaluated using metrics sensitive to both their magnitude at each timepoint and their overall shape across time. The impacts of undesired sensory potentials resulting from TMS pulse and muscle contractions were also assessed on both measures. Both conditioned-MEPs and TEPs were sensitive to re-afferent somatosensory activity following motor-evoked responses, but over different post-stimulus timepoints. Moreover, the amplitude of low-frequency oscillations in TEPs was strongly correlated with the sensory potentials, whereas early and local high-frequency responses showed minimal relationships. Accordingly, conditioned-MEPs did not correlate with TEPs in the time domain but showed high shape similarity with the amplitude of high-frequency oscillations in TEPs. Therefore, despite the effects of sensory confounds, the TEP and MEP measures share a response component, suggesting that they index a similar cortical response and perhaps the same neuronal populations.

## 1. Introduction

Transcranial magnetic stimulation (TMS) is a non-invasive brain stimulation method capable of activating cortical neurons across the intact scalp in humans [1]. A single TMS pulse depolarises a combination of excitatory and inhibitory neurons in the underlying cortical tissue, resulting in fluctuating periods of net excitation and inhibition that can last for several hundred milliseconds following stimulation [1]. In addition to local cortical circuits, TMS also indirectly activates cortical regions that are structurally connected to the stimulated region, resulting in a complex cascade of neural firing through large-scale cortical networks [2]. As such, TMS has emerged as a powerful tool to study excitatory and inhibitory neurotransmission in local and connected brain regions in humans.

The local and remote effects of TMS are typically quantified using some observable output. In the most common method, TMS of the primary motor cortex (M1) is coupled with electromyographic (EMG) recordings of a peripheral muscle targeted by the stimulated region. A single TMS pulse to M1 can result in a measurable muscle response known as a motor-evoked potential (MEP), the amplitude of which provides a measure of corticomotoneuronal excitability at the time of stimulation [3]. Paired-pulse TMS (ppTMS)-EMG paradigms are used to probe TMS-evoked changes in net excitation of motor-related cortical circuits across time, although the excitability of spinal circuits may also contribute [1, 4–6]. In these paradigms, a conditioning stimulus is delivered over M1 followed by a test stimulus (TS) after a given interval. The mean amplitude of the resulting conditioned MEP is then compared against the mean amplitude of the MEP induced by a TS alone to assess the excitatory or inhibitory effects of the conditioning stimulus [1, 7, 8]. In addition to local assessments of the stimulated M1, ppTMS-EMG can probe the interactions of M1 with other cortical regions by employing a second coil and applying the conditioning stimulus to an interconnected brain region (e.g., contralateral M1) [9]. Although ppTMS-EMG provides a wealth of information about local and inter-regional cortical circuits, this technique is limited to the assessments of motor circuits and provides an indirect measure of cortical activity which can be influenced by spinal excitability.

The second and more recent method combines TMS with electroencephalography (EEG), which measures fluctuations in cortical excitability across the scalp. The brain’s response to single-pulse TMS (spTMS) appears as a sequence of reproducible voltage deflections known as TMS-evoked EEG potentials (TEPs) which last up to 500 ms post-stimulus, at the site of stimulation and in interconnected brain regions [2, 10, 11]. M1 TEPs have been commonly characterized by seven peaks: N15, P30, N45, P60, N100, P180, and N280 [12, 13]. In general, EEG deflections such as TEPs are thought to reflect synchronous changes in postsynaptic potentials across large populations of neurons. Therefore, similar to ppTMSMEPs, changes in their amplitude can be used to quantify fluctuations in intracortical excitability following a TMS pulse, but with less influence from spinal circuits. However, in contrast to MEPs, the polarity of TEPs does not provide a clear indication of the net excitation/inhibition of cortical circuits (see Ref. [14] for a comprehensive explanation of polarity in EEG). Furthermore, some TEP components are either generated or modulated by peripherally-evoked potentials (PEPs) (e.g., auditory evoked potentials from the click and somatosensory evoked potentials from coil vibration or activation of scalp muscles [15, 16]) and/or proprioceptive sensory feedback (from the contraction in target muscles induced by suprathreshold stimulations of motor cortex) [17, 18]. Nonetheless, a growing body of evidence from pharmacological [19, 20] and behavioural TMS-EEG studies has linked certain peaks of M1 TEPs to periods of net excitation (e.g., N15, P30) and inhibition (e.g., N45, N100) [21, 22]. For instance, the reduction in MEP amplitude following ppTMS with an ISI of 100 ms (known as long-interval cortical inhibition) correlates with the size/shape of the N100 measured using EEG following TMS [22–24], suggesting the two measures may reflect a common inhibitory mechanism.

In addition to the time domain, TMS-EEG measures of cortical excitability can be examined in the frequency domain. Spectral decomposition of TMS-evoked EEG signals allows studying of oscillatory features (amplitude/power and phase) at various frequencies each reflecting distinct neurophysiological mechanisms [25]. Several studies have reported a transient increase in the power of ongoing alpha-band (8–12 Hz) and beta-band (13–30 Hz) oscillations in response to spTMS over motor cortex [26–28]. Furthermore, spontaneous fluctuations in the power [29–31] and phase [32, 33] of alpha oscillations, as well as the power of beta oscillations [21, 34], have been linked with changes in MEPs amplitude suggesting the two measures may reflect common changes in motor cortical excitability. However, these findings have not always replicated [29] and it is unclear whether TMS-evoked oscillations show similar relationships with MEP amplitude to spontaneous oscillations.

Taken together, both ppTMS-EMG and spTMS-EEG measure periods of net excitation and inhibition across time in response to a TMS pulse. What remains unclear is whether the responses measured by the two techniques index the same cortical activity. The aims of this study were two-fold: 1) to further explore the relationship between EEG (temporal and spectral) and EMG measures of TMS-evoked responses across time, using metrics sensitive to both the amplitude and shape of the signals [29]; and 2) to assess the contribution of peripheral sensory activity following the TMS pulse on the evoked responses measured in the frequency domain and with MEPs. We conducted two experiments and examined both local and interhemispheric responses to TMS over M1 at different intensities: sub- and supra-threshold. We also assessed the generalizability of the results by examining different stimulation waveforms (biphasic and monophasic) and evaluated the specificity of the findings to cortical stimulation by comparison with a sensory control condition (shoulder stimulation). Our primary hypothesis was that the amplitude of conditioned MEPs and TEPs measured from scalp electrodes over the M1 stimulation site would correlate across different post-stimulus timepoints, resulting in a similar shape between the two signals. Additionally, we explored how the amplitude of conditioned MEPs correlates with the magnitude of TMS-evoked oscillations at different frequency ranges.

## 2. Methods

### 2.1. Participants

Two experiments were conducted for this study. In Experiment I, 20 (24.50 ± 5 years; 14 females) right-handed healthy individuals were examined and in Experiment II, 16 (25 ± 6 years; 11 females) individuals were examined. Twelve subjects participated in both experiments. All subjects were screened for any contraindications to TMS [35], and provided written consent to the experimental procedure, which was approved by the Monash University Human Ethics Committee in accordance with the declaration of Helsinki. The TMS-EEG data from this study were also used in our recent publication investigating sensory contributions to TEPs [15]. Participants were seated comfortably in an adjustable chair, with their elbows resting on the armrests and forearms pronated and supported on a pillow on their laps. They were instructed to keep their eyes open, look at a black screen in front of them and stay relaxed.

### 2.2. TMS

In Experiment I, biphasic TMS (anterior-posterior/posterior-anterior) pulses were administered to the left M1 using a figure-of-eight coil (C-B60) connected to a MagPro X100+Option stimulator (MagVenture, Denmark). Stimulations were applied over the optimal point for the stimulation of first dorsal interosseous (FDI), where suprathreshold pulses consistently produced largest MEPs [36]. This location was digitally marked on each individual’s T1 scan using a neuronavigation system (Brainsight™ 2, Rogue Research Inc., Canada) to ensure the consistency of coil positioning across stimulation blocks. Resting motor threshold (rMT) was determined as the minimum stimulation intensity required to evoke MEPs greater than 50 μV in at least 5 of 10 successive trials [3]. Then, the intensity was gradually increased until MEPs of ~1 mV were elicited in at least 10 consecutive trials (S1mV). All assessments and stimulation blocks were carried out with an EEG cap on and the intensities are expressed as a percentage of maximum stimulator output (% MSO). Paired-pulse stimulations were delivered using either a sub- or suprathreshold conditioning stimulus (80% or 120% rMT) combined with a suprathreshold test stimulus (S1mV), at eight inter-stimulus intervals (ISIs): 15, 30, 45, 55, 100, 120, 180, and 220 ms. For MEPs, each subject received nine blocks of 40 stimuli over left M1. This included one block of 40 single-pulse stimuli at S1mV (spTMS, unconditioned), and eight blocks for which 40 paired-pulse stimuli were delivered at a given ISI (ppTMS, conditioned) with 20 pulses at each conditioning intensity. The stimulation intensities within each block were controlled by the MATLAB-based MAGIC (MAGnetic stimulator Interface Controller) Toolbox [37]. To examine TEPs, participants received four blocks of 50 spTMS each with two intensities of 80% and 120% rMT (100 pulses in total for each intensity), while EEG was continuously recorded.

In Experiment II, both uni- and bi-hemispheric stimulations were applied. TMS involved monophasic pulses in posterior-anterior direction delivered via Magstim 200^2^ stimulators (Magstim Company, UK) interfaced with a BiStim^2^ timing module (Magstim Company, UK). The stimulators were triggered by Signal software (V6) and CED data acquisition interface (Cambridge Electronic Design, Cambridge, UK). Uni-hemispheric stimulation was delivered to the left M1 through a figure-of-eight D70^2^ Coil (Magstim Company Ltd., Whitland, UK), and for bi-hemispheric stimulations a smaller second coil (D50 Alpha; Magstim Company Ltd., Whitland, UK) was positioned over the right M1. Using a smaller coil allowed positioning of both coils over the head. All of the baseline measurements and intensity settings were performed for each coil separately. Paired-pulse stimulations were delivered using suprathreshold conditioning stimulus (120% rMT) and TS (S1mV) with eight ISIs: 10, 20, 30, 40, 50, 100, 150, and 200 ms. For uni-hemispheric tests, TMS was applied over the left M1 and MEPs were recorded from right FDI. For bi-hemispheric TMS, the conditioning stimulus was also applied to the left M1 with the TS (S1mV) administered to right M1 while MEPs were recorded from left FDI. This method allowed us to examine how the activation of left M1 influenced the excitability of right M1. For MEPs, each individual received five blocks of uni-hemispheric and five blocks of bi-hemispheric TMS. Each block consisted of 40 ppTMS (5 for each ISI) and 10 spTMS (S1mV), pseudo-randomly distributed. For TEPs, participants received 75 single pulses over left M1 at 120% rMT while EEG was continuously acquired. Regular breaks were given between blocks. The interval between TMS pulses was jittered between 4 and 6 s.

For both experiments, white noise was played to the participants through earphones and a thin layer of foam was attached underneath the coil during all of the stimulation blocks [38] to attenuate the effect of PEPs on TEPs. Additionally, as a control condition, each participant received 100 suprathreshold TMS pulses over the left shoulder (around the acromioclavicular joint) to elicit PEPs due to TMS coil clicks and tapping sensation, without transcranially stimulating the brain [15]. While this control is likely suboptimal for somatosensory activation (e.g. shoulder vs scalp), we have previously demonstrated that the resulting sensory signal accounts for much of the fronto-central N100/P180 complex observed in motor TEPs using this data [15, 39]. To assess whether peripheral sensory input from the conditioning pulse could impact motor cortical excitability in ppTMS-EMG measures, we applied a conditioning stimulus over left shoulder and test stimulus over left M1 at ISIs of 100 and 200 ms (when the peaks of PEPs occur) in a subset of six participants.

### 2.3. EMG

EMG was recorded from the left and right FDI using bipolar Ag-AgCl surface electrodes (~2 cm apart) positioned in a belly-tendon montage. A ground electrode was fixed on the dorsum of the hand over the midpoint of the third metacarpal bone for each side. EMG was band-passed filtered at 10 to 1000 Hz, amplified 1000 times, sampled at 5 kHz, epoched around the stimulation pulse (–200 to 500 ms), and recorded on a computer for offline analyses. EMG data was processed offline using Labchart 8 (ADInstruments, Sydney, Australia). First, the trials with pre-stimulus muscle contractions (up to 100 ms) were identified and excluded following visual inspection. Then, the average of peak-to-peak values of MEPs evoked by each type of stimulation were calculated for each subject. Finally, the strength of inhibition or excitation was calculated as the percentage of the mean MEPs evoked by conditioned pulses (ppTMS) over the mean MEPs evoked by unconditioned pulses (spTMS, S1mV).

### 2.4. EEG

EEG was continuously acquired through a TMS-compatible EEG system (SynAmps^2^, Neuroscan, Compumedics, Australia) using 62 Ag/AgCl ring electrodes, arranged in an elastic cap according to the standard 10–20 layout (EASYCAP, Germany). The positions of the electrode were digitized and co-registered to each individual’s T1 weighted MR scan in the neuronavigation system. All the electrodes were grounded to AFz and online-referenced to FCz. EEG signals were amplified (1000 ×), low-pass filtered at DC–2000 Hz and digitized with a sampling rate of 10 kHz. The skin-electrode impedance level was kept below 5 kΩ throughout the recordings [12]. The signals were displayed and stored on a computer using the Curry8 software (Neuroscan, Compumedics, Australia). EEG data were pre-processed based on the method described in [40, 41], performed by custom scripts written in MATLAB (R2016b, The Mathworks, USA) using the functions provided by EEGLAB [42] and TESA [40] toolboxes. The cleaning pipeline has been described in detail in our recent publication [15].

To reduce volume conduction, EEG data was spatially filtered using a reference-free spherical spline surface Laplacian (spline flexibility = 4, spline regularization = 10^-5^) [43] and transformed into current source density estimates [44]. In addition to the time domain, TEPs were also assessed in the frequency domain. To extract the frequency and amplitude of TMS-evoked oscillations over time, Morlet Wavelet decomposition was performed for the average of trials [45] (width: 3.5 cycles; time steps: 1 ms; frequency steps: 1 Hz). The frequency range between 8 and 45 Hz was assessed to cover the dominant oscillations of the sensorimotor cortex [46, 47]). Subsequently, the oscillations in each frequency bin were normalised to baseline by dividing the post stimulus power by pre-stimulus power (averaged between -600 and -100 ms). Next, the frequency ranges with the strongest power following the TMS pulse were identified and a zero-phase Butterworth filter (4th order) was used to filter the time domain EEG data within these frequency ranges (8–12Hz and 13–45Hz). Finally, a Hilbert transformation [48] was performed on the filtered signal to estimate the instantaneous amplitude (envelope) of each frequency band across time.

To isolate left and right sensorimotor potentials, we extracted data at C3 and C4 electrodes, which approximates the left and right motor cortices respectively in the 10-20 system [49, 50], and grouped them with their eight neighbouring electrodes. As such, the average of potentials recorded at C3, FC1, CP1, FC5, CP5, C1, FC3, CP3, and C5 was calculated to represent the activity of left M1, and the average of the recordings at C4, FC2, CP2, FC6, CP6, C2, FC4, CP4, and C6 was taken as the activity of right M1.

### 2.5. Data analysis

#### 2.5.1 Comparing the oscillatory amplitude of TEPs and PEPs

All statistical analyses were performed in MATLAB. According to our previous findings [15], long latency TEPs (> 60 ms) recorded at the electrodes over the contralateral hemisphere can be highly contaminated by TMS-induced PEPs, despite adopting sensory attenuation measures (e.g., white noise and foam padding). However, the impact of PEPs on TMS-evoked oscillations has yet to be characterised. Hence, for each individual, we evaluated correlations between the magnitude of oscillations in spTMS-EEG and PEP signals using Spearman’s correlation coefficients, in both spatial (between topographic distribution of potentials at each time point) and temporal (between the recordings of the same electrodes across time) domains. For group-level analyses, we transformed the correlation coefficients (ρ) to *z* using Fisher’s transformation and applied one-sample permutation tests (10 000 iterations with *t*_max_ correction) to examine the *z*-scores against zero at each timepoint [51, 52]. The *z* scores were then transformed back to ρ values for illustration purposes. Previously, we showed that a spatial filter called signal-space projection with source-informed reconstruction (SSP–SIR) can effectively attenuate PEP contaminations in TMS-EEG signals [15, 53]. Therefore, where a relationship was found between the two measures, we repeated the assessment following filtering TEPs through SSP–SIR to rule out the confounding effect of sensory potentials.

#### 2.5.2 Comparing the effect of motor response size on TEPs and conditioned MEPs

Several studies have shown a link between the size of MEPs and temporal or spectral features of EEG recordings following motor cortex stimulations [17, 18]. The modulation of the TEP is thought to result from reafferent somatosensory activity following movement of the finger related to the MEP. To evaluate whether a similar phenomenon was present in our data, we split the spTMS-EEG trials into two conditions based on the peak-to-peak amplitude of their induced MEPs: high-MEP and low-MEP conditions containing the trials that evoked MEPs larger and smaller than the median of all MEPs; respectively. Then we compared the voltage levels of TEPs between the two conditions, across time and space, using cluster-based permutation tests as implemented in FieldTrip toolbox [54]. To assess whether conditioning MEP size influences the amplitude of the subsequent test MEP, we took a similar approach and split ppTMS-EMG trials based on MEPs induced by CS. Then we compared the amplitude of MEPs induced by TS between the two conditions using a permutation-based paired *t*-test with *t*_max_ correction [55].

#### 2.5.3 Comparing conditioned MEPs and TEPs

To examine the effect of different conditioning stimuli (80% and 120% rMT) on the size of MEPs induced by TS, we applied nonparametric Wilcoxon signed rank tests (as MEPs were not normally distributed at all timepoints) comparing the size of conditioned MEPs (induced by ppTMS) with unconditioned MEPs (induced by TS alone) at each ISI. The *p*-values were then corrected for multiple hypothesis testing by controlling the false discovery rate (FDR) using the method of Benjamini Hochberg [56]. To assess the relationship between EEG and EMG measures of cortical responses to TMS, we compared conditioned MEP amplitude at each intensity of conditioning stimulus with the amplitude of EEG potentials evoked by the conditioning stimulus alone (spTMS), adopting two main approaches. First, we examined Spearman’s correlations between the two types of measures at each point of time (each ISI) to see whether the amplitude of EEG potentials is related to the net excitation or inhibition observed in MEPs. Second, we compared the shape of the two signals (patterns of fluctuations, independent of amplitude) using a shape-similarity metric (SSM) (Fig. 1). The SSM between MEPs and EEG measures was calculated at the single subject level as the sum of the absolute differences between the MEPs and EEG measures over all ISI time points, allowing an arbitrary proportional rescaling on EEG signals:

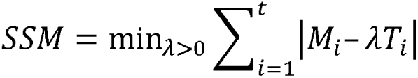

for a rescaling parameter, λ, across *t* time points, where *M* is MEP measurement, and *T* is EEG measurement (amplitude of TEP or instantaneous amplitude of oscillations). λ was first constrained to positive (i.e., no polarity change in EEG) and then negative (reversing polarity in EEG) values, to assess direct and inverse relationships between the two signals, separately. The SSM metric is formulated to be invariant to a proportional rescaling of either *M* or *T*, and therefore captures the similarity in relative variation (shape) of the two signals, with lower SSM values indicating that *M* and *T* have similar shapes. Conceptually, measuring SSM is similar to performing a correlation analysis, but has the advantage of maintaining the baseline zero for *M* and *T*, and controlling (preserving or reversing) the polarity of the signals (via the constraint λ > 0 or < 0). To examine whether a computed SSM value was significantly lower than what would be expected by chance, we randomly shuffled EEG and EMG data to compute a 1000-sample null distribution of SSM values. This null distribution was used to compute *p*-values for SSM directly, as a permutation test. For the group-level analysis, we calculated the sum of SSM values across all individuals and evaluated the statistical significance as a permutation test, using the null distribution formed by sum of the SSM values from the individually permuted EEG and EMG data (Fig. 1). EEG signals before 15 ms (ISI = 10 ms) are excluded for this analysis, due to the high level of contaminations with TMS-induced artefacts.

**Figure 1:**
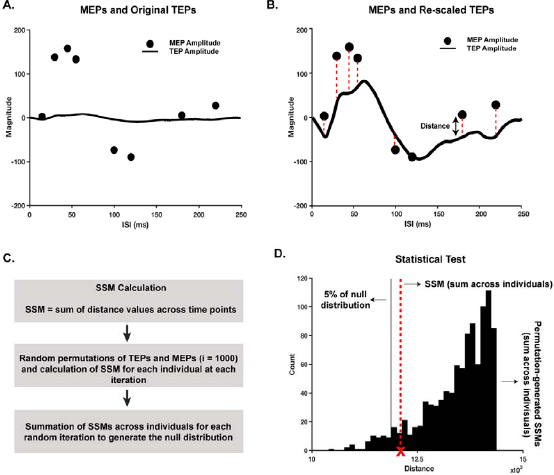
Calculation and statistical evaluation of shape-similarity between MEPs and TEPs. MEPs represent the amplitude of conditioned relative to unconditioned MEPs (%) and TEPs represent the amplitude of the EEG potentials (μV). In this example, the shape similarity between TEPs and MEPs is not statistically significant (D), as the result SSM is within the range of 95% of the permutation generated SSMs.

### 2.6 Data and code availability

The pre-processed EMG and EEG data are freely available on figshare at: https://doi.org/10.26180/5cff2a0fc38e9. Raw data are available upon request. The code for processing the EEG data is available at: https://github.com/BMHLab/TEPs-PEPs. The code for comparing EEG and EMG output measures is available at: https://github.com/BMHLab/TEPs-MEPs.

## 3. Results

The study protocol was well-tolerated by all participants and no adverse effects were reported. In experiment I (MagVenture, Biphasic), the number of spEEG trials following cleaning were 78.0 ± 12.0 and 82.4 ± 9.5 (average across individuals) for supra- and subthreshold responses; respectively. RMT was 57 ± 9 % MSO and the amplitude for S1mV was 70 ± 12% MSO. In experiment II (Magstim Bistim, Monophasic), 69.5 ± 23.5 spEEG trials were preserved following cleaning. rMT and S1mV were 48 ± 7 and 60 ± 8 % MSO for left M1 (D70 coil), and 58 ± 5 and 72 ± 8 % MSO for right M1 (D50 Alpha coil), respectively. Table S1 presents the number of clean ppTMS-EMG trials at each ISI.

### 3.1. EMG and EEG responses to TMS over M1 and shoulder

Following suprathreshold biphasic stimulation over left M1 (conditioning stimulus at 120% rMT), conditioned MEPs exhibited an early phase of facilitation (ISIs < 30 ms) followed by a long phase of inhibition (ISIs ~60 to 180 ms) (Fig. 2A). M1 TEPs evoked by spTMS (at the same intensity as conditioning stimulus in ppTMS), showed a sequence of strong deflections with an early negativity around 20 ms followed by a period of positivity (from ~30 to 80 ms) and a long period of negativity (from ~80 to 220 ms; Fig. 2B). In the frequency domain, TMS evoked two main oscillatory patterns at a higher frequency (~13–45 Hz) peaking around 30 ms, and a lower frequency oscillation (~8–12 Hz) peaking at ~100 ms (Figs 2C-F). Both ppTMS-EMG and spTMS-EEG following suprathreshold monophasic stimulation showed similar patterns as observed in the biphasic condition (Fig. S1).

**Figure 2:**
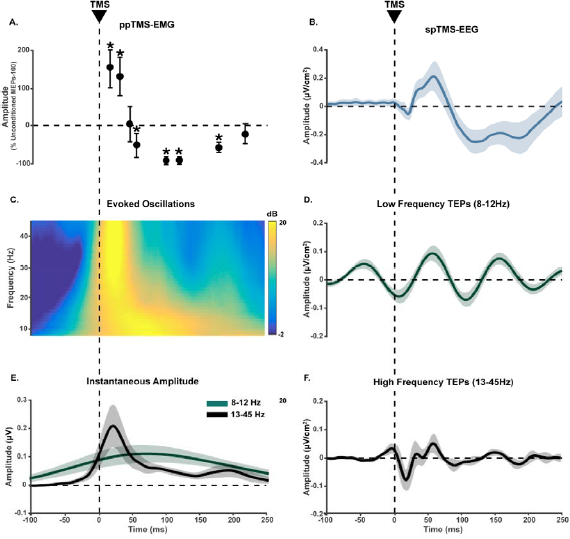
EMG and EEG measures of cortical responses to suprathreshold biphasic TMS. A) ppTMS-EMG recordings. Both conditioning stimulus and test stimulus were administered to left M1 and MEPs were recorded from right FDI. The solid circles represent the mean amplitude of conditioned MEPs expressed as a percentage of the mean unconditioned MEP amplitude (across individuals) from TS alone. The results are subtracted by 100 to show inhibition as a negative value. The error bars indicate 95% of confidence intervals assuming a Gaussian distribution. *indicate *p* < 0.05 comparing conditioned and unconditioned MEPs. B) spTMS-EEG recordings in the time domain. TMS is applied over left M1 with the same intensity as the conditioning stimulus in ppTMS. TEPs show the average of recordings at C3, FC1, CP1, FC5, CP5, C1, FC3, CP3, C5. The shaded areas show the 95% of confidence intervals. C) Distribution of the evoked oscillatory powers of different frequency bins recorded at left M1 across time, obtained from Morlet wavelet decomposition. D-F) TEPs filtered into the frequency bands with maximum power (observed in C). E) Instantaneous amplitude of low and high frequency oscillations. The line graphs show the group average of Hilbert amplitude changes recorded at left M1 and the shaded areas represent the 95% of confidence intervals.

Following subthreshold biphasic stimulation, the magnitude of MEPs to conditioned TS did not statistically differ from the MEPs to TS alone, at any of the ISIs (*p* > 0.05), although a small facilitation was detectable at 15 ms, consistent with intracortical facilitation. Comparably, the induced TEPs only showed small early deflections prior to 100 ms post-stimulus. Frequency analysis showed similar oscillatory patterns as observed in suprathreshold condition, although with lower power levels (Fig. S2).

For inter-hemispheric stimulation, conditioning stimulus over left M1 reduced the MEPs following TS over right M1 at ISIs of 10 to 50 ms, corresponding to the two phases of IHI [9](Fig S3). TEPs recorded at right M1 (following left M1 stimulation) showed an early positive peak around 30 ms, followed by a period of negativity from 40 to 130 ms and then a large positive peak around 180 ms. Frequency analysis showed similar oscillatory patterns as observed in local responses (early fast followed by slow oscillations), although with lower power levels.

EEG signals following shoulder stimulation (control condition; PEPs) showed small early fluctuations followed by two large peaks at around 100 and 200 ms (Fig. 3A). In the frequency domain, slow oscillations were dominant following shoulder TMS with maximum power between 100 and 200 ms (Fig. 3B). High-frequency oscillations were mainly observable at earlier responses and, in total, had lower magnitude relative to the slow oscillations (Fig. 3C).

**Figure 3:**
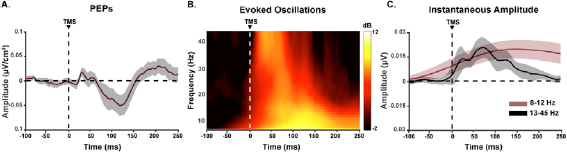
EEG measures of cortical responses to spTMS over left shoulder. A) Changes in PEPs amplitude recorded at left M1 (C3, FC1, CP1, FC5, CP5, C1, FC3, CP3, C5) in the time domain. B) Distribution of the evoked oscillatory powers of different frequency bins recorded at left M1 across time, obtained from Morlet wavelet decomposition. C) Group average changes of Hilbert amplitude in left M1 recordings. Measures in B and C are baseline-corrected (−600 to -100ms). The shaded areas in A and C represent the 95% of confidence intervals.

### 3.2. Relationships between spectral features of TEPs and PEPs

In a previous study, we showed high correlations between TEP and PEP signals in the time domain following motor cortex stimulation [15], however the relationship between TEPs and PEPs in the frequency domain has not been reported. As such, we first compared both high- and low-frequency oscillations following cortical and shoulder (control) stimulation. Following cortical TMS to motor cortex, maximum oscillatory power centred over sensorimotor electrodes for both high- and low-frequency oscillations. In contrast, the strongest oscillations following shoulder stimulation were recorded at central electrodes (Fig. 4A). Moreover, comparisons of the oscillatory magnitudes revealed that TMS over M1 evokes stronger low-frequency oscillations at the side of stimulation and stronger high-frequency oscillation across the whole scalp (Fig. 4B). Nevertheless, Spearman’s correlation tests revealed that the two signals were highly correlated across certain temporal and spatial ranges. Specifically, for low-frequency oscillations, we found strong correlations between conditions over central and contralateral electrodes during the almost the whole post-stimulation time (Figs 4C-D). For high-frequency oscillations, strong correlations were observed over contralateral sensorimotor electrodes occurring between ~50 and ~160 ms (peaking before 100 ms; Fig. 4E). The results suggest that PEPs likely contribute to TMS-evoked oscillations following motor cortex stimulation, although not over the site of stimulation.

**Figure 4:**
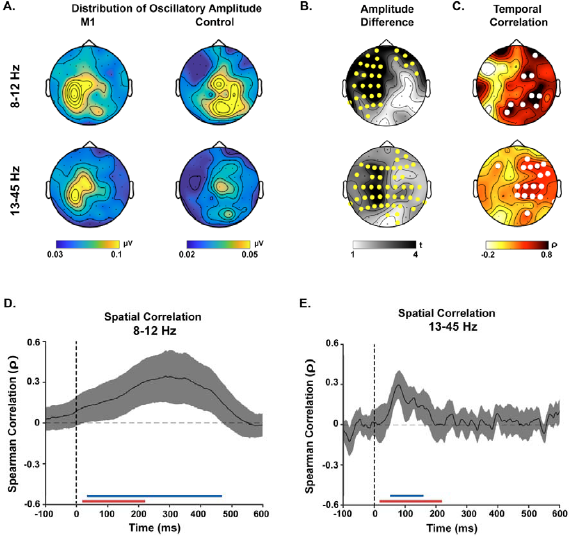
Comparisons of oscillatory amplitude between cortical and shoulder stimulation. A) Scalp maps illustrate the spatial distribution of instantaneous Hilbert amplitude at each condition. B) Results of cluster-based permutation tests comparing the magnitude of oscillations between conditions. The highlighted electrodes showed a significant difference between the conditions (*p* < 0.05). C) Distribution of temporal correlation of oscillatory amplitude between conditions. The highlighted electrodes showed significant correlation (*p* < 0.05). A-C) The scalp maps represent the time between 16 and 220 ms post-stimulus. D-E) The line graphs demonstrate changes in the spatial (topographical) correlations between conditions. The shaded areas represent the 95% of confidence intervals and the dash vertical lines show the moment of stimulation. The red line right above the *x*-axis highlights the time window used in this study to compare TMS-EMG and TMS-EEG potentials. The blue line (above the red line) represents the time when the two signals showed significant correlations in amplitude (*p* < 0.05).

### 3.3. Effects of motor response amplitude on conditioned MEPs and TEPs

To evaluate the effect of sensory inputs from muscle contractions resulting from MEPs on subsequent conditioned MEPs and TEPs, we split suprathreshold trials into low- and high-MEP conditions based on the amplitude of CS-induced MEPs. We did not consider ISI 15 ms for this analysis as the responses to CS and TS were not reliably separable. The average of TS-induced MEP amplitude at each ISI is shown in Figure 5A. Compared to low-MEP trials, conditioned MEPs were larger in high-MEP trials at almost all ISIs, although the difference was statistically significant only at 30 ms (paired-sample permutation test with *t*_max_ correction for multiple comparisons, *p* < 0.0001). For TEPs, the overall amplitude, as measured by the global mean field power, was significantly stronger in high-MEP trials compared to low-MEP trials (paired-sample permutation test with *t*_max_ correction for multiple comparisons, *p* = 0.001). Cluster-based permutation tests including a time-range between 15 and 220 ms revealed that this difference was strongest in TEPs from sensorimotor electrodes between ~45 and ~100 ms and at ~220 ms post-stimulus (Fig. 5C). We could not find any strong evidence that conditioning MEP amplitude impacted the magnitude of either low- (Fig. 5D) or high- (Fig. 5E) frequency oscillations in TEPs. Together, these findings show that the amplitude of conditioned MEPs and TEPs are both impacted by the size of the initial MEP, although over different time periods.

**Figure 5:**
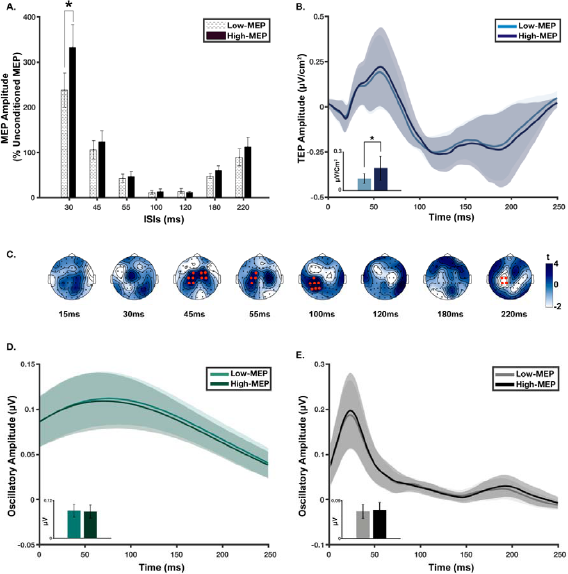
Effect of MEPs amplitude on EMG and EEG measures of cortical responses to suprathreshold and biphasic TMS. A) Amplitude of TS-induced MEPs at each ISI. The error bars represent the standard error of measurements (SEM). *indicates statistically significant differences (corrected *p* < 0.0001). B) Comparison of TEPs amplitude between high and low-MEP conditions. *indicates statistically significant difference (*p* = 0.004). C) The scalp maps compare the voltage distribution of TEPs from the two conditions, at each ISI, using cluster-based permutation tests. The channels highlighted in red represent the clusters that showed statistically stronger responses in high-MEP condition. One positive and one negative significant cluster were found (Monte Carlo corrected *p* = 0.0001 and 0.014; respectively). C-D) Comparisons of the oscillatory amplitude of EEG potentials between high and low-MEP conditions. B, D-E) The solid lines represent the group-averaged of TEPs (B), Hilbert amplitude at 13–45 Hz (D) and 8–12 Hz (E) obtained from EEG signals recorded at left M1 (C3, FC1, CP1, FC5, CP5, C1, FC3, CP3, C5). The shaded areas show the 95% of confidence intervals (CI). All oscillatory measures are baseline-corrected (−600 to -100 ms). The embedded bar plots compare the average of the absolute values of each measure overt time (15-250 ms).

### 3.4. Correlations between conditioned MEPs and TEPs at different time points

We aimed to examine whether the level of cortical excitability measured by different ppTMS-EMG paradigms (with different ISIs and conditioning stimulus intensities) correspond to the magnitude of TMS-EEG potentials. Hence, we correlated the amplitude of conditioned MEPs with the amplitude of TEPs (in both time and frequency domains) evoked by spTMS with the same intensity as conditioning stimulus and at the corresponding timepoint to the ISIs. In general, conditioned MEPs following suprathreshold stimulation showed weak correlations with TEPs at all evaluated timepoints. The strongest relationship was found between conditioned MEPs and low frequency amplitude at 220 ms (*ρ* = 0.50; *p*_Uncorr_ = 0.03), although the correlation did not remain statistically significant after correcting for multiple tests (p_corr_ = 0.21). Similarly, there was no strong evidence for relationships between EEG and EMG responses to suprathreshold monophasic stimulation (Fig. 6A).

**Figure 6:**
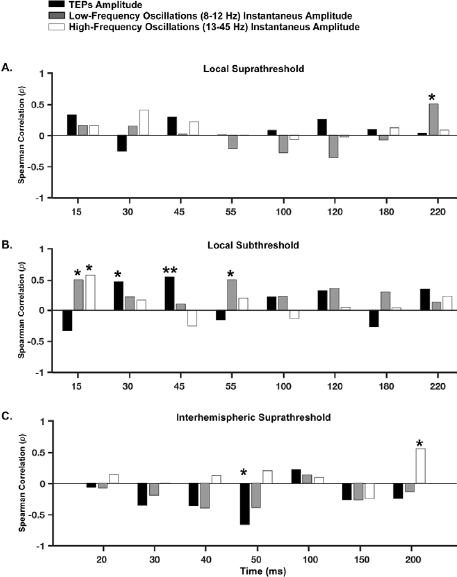
Spearman correlations between the amplitude of conditioned MEPs and TEPs at each point of time. A) Local responses to biphasic and suprathreshold TMS. B) Local responses to biphasic and subthreshold TMS. C) Interhemispheric responses to monophasic and suprathreshold TMS. Time axis represents the ISIs in ppTMS; the time points at which spTMS-EEG recordings are extracted for correlation analysis. * represents *p*_uncorr_ < 0.05. **indicates p_corr_ < 0.05.

In the subthreshold condition, conditioned MEP amplitudes showed strong relationships with TEPs at earlier time points (Fig. 6B). The strongest correlations were found between conditioned MEPs and TEP amplitudes at an ISI of 45 ms (ρ = 0.55; *p*_corr_ = 0.1) and high frequency power at 15 ms (ρ = 0.58; *p*_corr_ = 0.07). In the inter-hemispheric condition, conditioned MEPs from the contralateral hemisphere (recorded from left FDI) and contralateral TEP amplitudes (recorded from right M1) gradually developed negative correlations over time peaking at 50 ms post-stimulus (ρ = -0.66; *p*_corr_ = 0.05) (Fig. 6C). Correlations between the amplitude of conditioned MEPs and high- and low-frequency oscillations were weak at all time points (p_corr_ > 0.05). Together, the findings provide moderate support for a relationship between the magnitude of early responses to subthreshold TMS across methods. However, there was no evidence for a relationship between the magnitude of MEP facilitation/inhibition and TEP amplitude following suprathreshold stimulation at the site of stimulation. For all of the evaluated conditions, we also split the high-frequency range into the canonical beta (13–30 Hz) and gamma (31–45 Hz) frequency bands but did not find any associations with conditioned MEPs (*p*_corr_ > 0.05))

### 3.5. Shape similarity between conditioned MEPs and TEPs

Next, we examined whether the two techniques (ppTMS-EMG and spTMS-EEG) measure the same pattern of changes in cortical excitability following a single TMS pulse overtime. Therefore, instead of focusing on the magnitude of potentials at individual points in time, we compared their overall shape of fluctuations across post-stimulus time. Following suprathreshold biphasic stimulation, we could not find evidence for similarities in shape between conditioned MEPs and TEPs (*p* = 0.12; positive relationship) or low-frequency oscillatory magnitude (*p* = 0.66; negative relationship) (Fig. 7A). However, fluctuations in high-frequency oscillatory amplitude were similar in shape with conditioned MEPs across time (*p* = 0.001). This relationship was found at the individual level in 11/20 participants and was also observed following monophasic pulses (*p* = 0.001; Fig. S4), and at subthreshold intensities (*p* = 0.05; Fig. 7B). In contrast, we could not find evidence for a similarity in shape between conditioned MEPs and TEPs in the interhemispheric condition (Fig. 7C). Splitting the high frequency oscillations into beta and gamma bands did not change the results in any of suprathreshold conditions (all *p* < 0.05). However, in subthreshold stimulation, only gamma oscillations remained similar in shape with conditioned MEPs (*p*(beta) = 0.1; *p*(gamma) = 0.03)). Together, the results suggest that the magnitude of high-frequency oscillations at the site of stimulation following TMS follow a similar time course (i.e., shape) to conditioned MEPs, even though the amplitude of these signals do not necessarily show a strong relationship at any given point in time.

**Figure 7:**
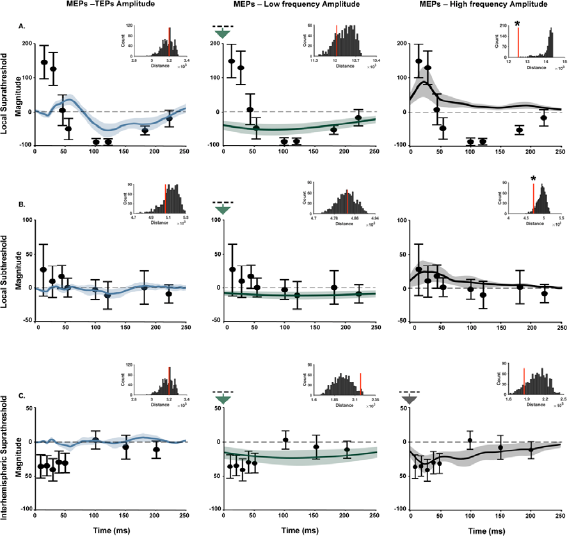
Shape similarity between ppTMS-EMG and spTMS-EEG measures of cortical excitability. A) Local responses to biphasic and suprathreshold stimulation over left M1. B) Local responses to biphasic and subthreshold stimulation over left M. C) Interhemispheric responses to monophasic and suprathreshold stimulation over left. For local responses (A-B), TEPs are averaged across C3, FC1, CP1, FC5, CP5, C1, FC3, CP3 and C5 electrodes, and MEPs are from right FDI. For interhemispheric responses (C), TEPs are averaged across C4, FC2, CP2, FC6, CP6, C2, FC4, CP4, and C6 electrodes, and MEPs are from left FDI. The solid circles represent the mean amplitude of conditioned MEPs expressed as a percentage of the group average of MEP amplitude from TS alone. The error bars indicate 95% of confidence intervals assuming a Gaussian distribution. The line graphs represent the group average changes in the amplitude of TEPs (left column), low-frequency (8–12 Hz; middle column) and high-frequency (13–45 Hz; right column) evoked oscillations, rescaled to the optimum rescaling values identified by SSM algorithm. The shaded areas show the 95% of confidence intervals. Instantaneous amplitude of oscillations is baseline-corrected (−600 to -100ms). The embedded bar plots depict the distribution of group sum of distances between randomly shuffled TEPs and MEPs (*x*-axis represents the measured distances and *y*-axis shows the frequency of each distance value). The red vertical line shows the group sum of the real distances. * shows that the real distance was smaller than 95% of the permutation-generated distances, suggesting that the similarity between the measures is statistically better than that would be expected by chance. The downward arrow on the top of *y*-axis indicates that EEG measures are rescaled to negative values.

### 3.6. Relationships between conditioned MEPs and control PEPs

Next, we asked whether any of the relationships we observed between conditioned MEPs and TEPs could be explained by PEPs. Spearman correlation tests did not provide any strong evidence for correlations between conditioned MEPs amplitudes and PEPs at any point in time (*p*_corr_ > 0.05). Furthermore, there was no evidence for similarities in shape between conditioned MEP and either PEP amplitude or high-frequency PEP power (Fig. 8). Interestingly, we did find evidence for shape similarity between conditioned MEPs and low-frequency PEP at the site of stimulation, which persisted across stimulation waveforms and intensities (Figs 8A-B and Fig. S4). In this relationship, the peak of MEP inhibition coincided with the peak in the magnitude of slow oscillations in EEG (around 100 ms), raising the possibility that peripheral sensory input could contribute to MEP inhibition in motor cortical circuits.

**Figure 8:**
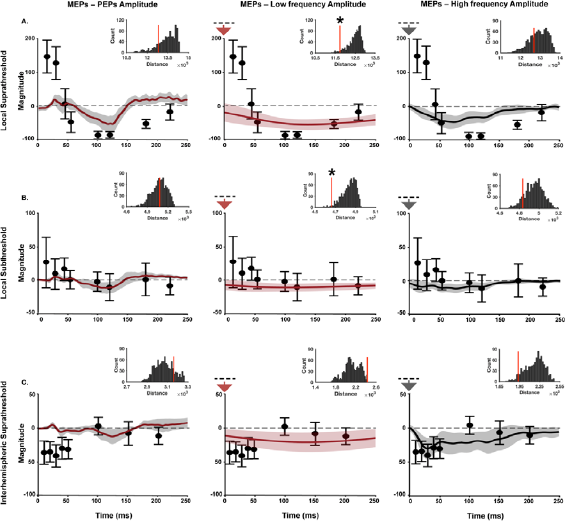
Shape similarity between PEPs and EMG measures of cortical excitability. A) local responses to suprathreshold and biphasic TMS. B) local responses to subthreshold and biphasic TMS. C) interhemispheric responses to suprathreshold and monophasic TMS. A-B) The line graphs represent group-averaged EEG recordings at left M1 (C3, FC1, CP1, FC5, CP5, C1, FC3, CP3 and C5) to biphasic spTMS over left shoulder. C) The line graphs represent group-averaged EEG recordings at right M1 (C4, FC2, CP2, FC6, CP6, C2, FC4, CP4, and C6) to monophasic TMS over left shoulder. The shaded areas show the 95% of confidence intervals. Oscillatory measures are baseline-corrected (−600 to -100 ms). The solid circles represent the group-averaged changes in MEPs (conditioned relative to unconditioned) recorded from right (A-B) and left (C) FDI in response to ppTMS over left M1(A-B) left-right M1s (C). The error bars indicate 95% of confidence intervals assuming a Gaussian distribution. The embedded bar plots depict the result of shape-similarity tests between the two signals. * shows significant similarity in shape between the two signals (*p* < 0.05). The downward arrow on the top of the *y*-axis indicates that EEG measures are rescaled to negative values.

In contrast to cortical stimulation, we found that applying the condition pulse to the shoulder were slightly stronger relative to unconditioned MEPs evoked by a test stimulus alone (tested in a subset of six participants; Fig. 9). These preliminary findings suggest that the strong inhibition of MEPs at 100 ms observed following cortical stimulation is unlikely the result of peripheral sensory input, and therefore the shape similarity between low-frequency PEPs and conditioned MEPs is likely spurious (i.e., the two measures share a similar time course but are unrelated).

**Figure 9:**
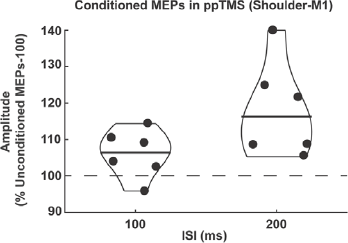
Changes in the amplitude of conditioned MEPs recorded at right FDI when the conditioning stimulus was applied over left shoulder. Black circles represent changes in MEPs for each individual and the horizontal solid lines indicate the median values.

Given that PEPs can result in potentially spurious relationships with conditioned MEPs, we reanalysed the relationship between conditioned MEPs and high-frequency TEPs. We used SSP-SIR to suppress PEPs in the TEP signal and then assessed whether the shape of conditioned MEP amplitudes was still related to the magnitude of fluctuations in high-frequency oscillations in TEP. Following PEP suppression, we still found evidence for shape similarity between conditioned MEPs and high-frequency oscillations following both suprathreshold (biphasic, *p* = 0.004; monophasic, *p* = 0.001) and subthreshold (*p* = 0.02) stimulation, suggesting the relationships between conditioned MEPs and high-frequency oscillations in EEG recordings are unlikely explained by PEPs.

## 4. Discussion

In this study we explored the relationship between EMG and EEG measures of TMS-evoked cortical excitation and inhibition both at the site of stimulation (left motor cortex) and in the contralateral hemisphere, using different stimulation parameters (intensity and waveform). We used two metrics (magnitude and shape) to compare MEP amplitude from ppTMS, with the amplitude and oscillatory manitude of spTMS-EEG recordings across time. There were three main findings. First, contrary to our hypothesis, we could not replicate the previous findings that showed a relationship between the N100 TEP amplitude/slope and inhibition of MEPs at ISIs of 100 ms following suprathreshold stimulation. In fact, we did not find any strong correlations between the magnitude of MEP and TEP measures at any point of post-stimulus time. Second, we found that the pattern of changes in the amplitude of conditioned MEPs over time was similar in shape with the fluctuations in the magnitude of high-frequency oscillations (~13–45 Hz) at the site of stimulation, regardless of stimulation intensity or waveform. The similarity in shape between signals could not be explained by general sensory activity following TMS (i.e., shoulder stimulation) and was not observed for lower-frequency oscillations. Third, we found that trials resulting in a larger MEP following the initial TMS pulse corresponded with larger conditioned MEPs at short ISIs (30 ms) and larger TEP amplitudes between 45 and 200 ms. The temporal decoupling of these effects suggests that EMG and EEG measures of TMS-evoked activity are differentially sensitive to peripheral factors, which may mask their relationship to cortical excitability. Together, these findings provide preliminary support that the early burst in power of high-frequency oscillations following TMS mirror the period of early facilitation in conditioned MEPs. Therefore, despite the effects of sensory confounds, the TEP and MEP measures share a response component, suggesting that they index a similar cortical response and perhaps the same neuronal population.

### 4.1. Relationship between conditioned MEPs and TEPs in the time domain

Several lines of evidence suggest that the N100 TEP peak represents a period of inhibition in the stimulated cortical region. For instance, the amplitude of this peak following M1 stimulation correlates with measures of motor cortex inhibition measured with EMG including the silent period [57] and LICI [22, 23]. The N100 TEP also increases in amplitude in response to cortical states associated with higher cortical inhibition, such as preparing to resist movement following a TMS pulse or wrist perturbation [58], and decreases in states associated with lower inhibition such as preparing to make a motor movement [59]. Furthermore, positive modulators of inhibitory GABA-B receptors, like Baclofen, significantly increased the amplitude of N100 providing pharmacological support that inhibitory neurotransmission at least partially underlies this component [19].

In the present study, we assessed whether: 1) we could replicate the relationship between the N100 peak amplitude and LICI; and 2) whether this relationship extended to other time periods, such as the early period of MEP facilitation. In contrast to previous studies, we did not find evidence suggesting a relationship between the amplitude of the N100 at rest and LICI at ISI of 100 ms. Furthermore, we could not find any relationship between TEP amplitudes and conditioned MEP amplitudes at any other time point following suprathreshold stimulation, and only moderate evidence for a relationship at 30 and 45 ms in the subthreshold condition (a period with little net excitation/inhibition). There are several possible interpretations of these findings. First, it is possible that the neural populations responsible for the broad-band TEP measured in the time-domain differ from those resulting in MEPs. MEPs are generated by transsynaptic excitation of a relatively small population of corticospinal pyramidal neurons targeting a specific muscle via connections with alpha motoneurons. In contrast, TEPs are sensitive to larger neural populations which synchronise activity from multiple cortical regions. The small population of neurons responsible for MEPs may not overlap with the populations of neurons generating the broad-band TEP signal, thereby decoupling the two measures.

Related to this point, both MEPs and TEPs are sensitive to different peripheral factors. In a previous study using the same data, we showed that the N100 following motor cortex stimulation represents at least two different sources: a fronto-central response which is common with sensory activity resulting from the TMS pulse (auditory and somatosensory inputs); and a left motor response which may represent transcranial cortical stimulation of the motor cortex [15]. In the current study, we have extended this finding by showing that the left motor N100 and earlier peaks including the N45 and P55 are modulated by the size of the resulting MEP. The MEP/TEP relationship replicates several previous findings, which also showed that TEP amplitudes are larger when the corresponding MEP is larger [17], and corresponds with increased connectivity from the primary sensory cortex to the primary motor cortex [45]. The increase in connectivity from sensory to motor cortices suggests the TEP modulation between high- and low-MEP trials may reflect differences in reafferent sensory activity following MEP-related movement of the fingers and hand. In contrast, conditioned MEP amplitude at ISIs of 30 ms, but not 100 ms, were modulated by the conditioning MEP size. The increase in conditioned MEPs at shorter ISIs with larger MEPs is consistent with increased spinal excitability resulting from the descending volley following the conditioning stimulus [60], but may also reflect modulation of spinal excitability by Ia afferents following muscle movement. The differences in sensitivity of TEPs and conditioned MEPs to peripheral factors across different time points may therefore mask any actual relationship between the aspects of the two measures sensitive to the excitability of the stimulated cortical region.

Finally, there were subtle differences in methodology with previous studies assessing N100/LICI relationships. For example, both [22] and [23] assessed the slope of the N100 evoked by the conditioning pulse in a paired-pulse paradigm as opposed to a separate single pulse condition as we did in this study. It is possible that slope and amplitude of TEPs may represent slightly different mechanisms. However, [23] only found a relationship between TEP slope and LICI at 100 ms intervals, but not 150 ms, partially in line with our findings. Taken together, our results suggest that conditioned MEPs and TEPs measured in the time domain are likely sensitive to different neural populations and different peripheral systems, and therefore provide differing, but complimentary measures of excitatory and inhibitory activity following TMS.

### 4.2. Relationship between conditioned MEPs and TEPs in the frequency domain

In addition to the time domain assessments, spectral decomposition of EEG recordings can reveal some distinct, yet complementary, aspects of neural responses following TMS [61]. However, the functional and physiological correlates of TMS-evoked EEG oscillations are largely unknown. The current results provided evidence for two distinct oscillatory responses to TMS over M1in both local and interhemispheric recordings: long-lasting low-frequency oscillations in the alpha range (~8–12 Hz) and short high-frequency oscillations in the beta/gamma range (~13–45 Hz) (Fig. 2C). Both rhythms showed some level of specificity to the site of stimulation during early timepoints; with maximum power at the sensorimotor areas and weak correlations with PEPs (Fig. 4). However, the two oscillatory ranges differed in their relationship with motor cortex excitability indexed by conditioned MEPs.

High-frequency oscillations are thought to represent a local cortical response to TMS over the motor cortex, but are also influenced by corticothalamic loops [27]. We found that the overall shape of high-frequency TEPs (within the range of beta and gamma frequencies) corresponded to the shape of changes in conditioned M1 across time, using different TMS waveforms (monophasic, biphasic) and intensities (subthreshold, suprathreshold). Furthermore, the instantaneous amplitude of high-frequency oscillations in TEPs correlated with MEP facilitation at 15 ms following a subthreshold conditioning pulse (i.e., intracortical facilitation), but not a suprathreshold conditioning pulse. Importantly, the finding could not be accounted for by sensory activation following a control condition (stimulation of the shoulder). While this control can rule out general sensory contributions to the signal, it does not mimic the scalp specific somatosensory input of the TMS pulse, which would require a separate control (i.e. electrical stimulation of the scalp). As such, we cannot rule out the possibility that somatosensory input from the scalp contributes to the high-frequency oscillations observed at the site of stimulation. Nonetheless, these results suggest that TMS-evoked high frequency oscillations may result from a similar neuronal population responsible for the facilitation of MEPs following a conditioning pulse.

Spontaneous changes in high-frequency oscillatory power are associated with modulation of corticospinal excitability. For example, higher pre-stimulus power, but not phase, in both beta and gamma-band oscillations are associated with larger MEPs [62]. These findings are broadly in line with our results, which suggest that the high-frequency oscillatory burst following a TMS pulse corresponds with a period of increased corticospinal excitability following both subthreshold and suprathreshold conditioning intensities. Our findings also suggest the power and not the phase of high-frequency oscillations are associated with increased corticospinal excitability, as MEPs are facilitated at ISIs of both 15 and 30 ms, periods that correspond with both negative (N15) and positive (P30) peaks of the high-frequency TMS-evoked burst. Interestingly, high-frequency TMS-evoked power did not track the strong inhibitory period corresponding with LICI, instead returning to pre-stimulus levels. It is possible that high-frequency oscillations do not represent active inhibition but are instead modulated by increased inhibitory tone. For example, early high-frequency TEP peaks including the P30 following the test pulse are also suppressed with paired pulse paradigms at ISIs of 100 ms [22, 23] and 150 ms [23] over motor cortex similar to MEPs. Furthermore, the early high-frequency peaks (N15, P30) from both motor and non-motor sites are modulated by sleep deprivation [63], repetitive TMS protocols [64], and drugs affecting voltage gated sodium channels [65], but not glutamatergic [66, 67] or GABAergic receptors [19, 67]. Together, these findings represent a growing body of evidence that the early high-frequency TEP peaks reflect a general measure of cortical excitability which is independent of glutamatergic and GABAergic neurotransmission.

For TMS-evoked alpha oscillations, we only found a weak (non-significant) negative correlation between the shape of changes in local magnitude and conditioned MEPs (Fig. 6). The relationship between alpha oscillations and corticospinal excitability is unclear. Some studies have found a negative correlation between pre-stimulus alpha power and the amplitude of MEPs [30, 68], while others have reported no associations [32, 69, 70]. This mixed evidence may suggest that spontaneous alpha oscillations in motor cortex do not strongly involve corticospinal neurons. Unexpectedly, we did observe a relationship between the shape of magnitude changes in PEP-evoked alpha oscillation and MEP inhibition (Figs 7A-B). However, we did not find evidence for MEP inhibition following shoulder stimulation (Fig 9), rendering it unlikely that MEP inhibition result from PEPs. The spurious relationship between PEPs and MEP inhibition highlights an important limitation of the correlational measures and the classical hypothesis tests used in the current study. More conservative statistical approaches that preserve the autocorrelations in signals might help achieve more robust results when correlating EMG and EEG potentials [71].

### 4.3. EMG and EEG measures of the interhemispheric response to TMS

Following a TMS pulse, activity is not restricted to the stimulated cortical region but also propagates to the interconnected areas. Previous ppTMS-EMG studies demonstrated two main periods of interhemispheric inhibition (IHI) between homologous motor cortices, peaking around 10 and 30 to 50 ms [9, 72, 73]. The present findings show both periods of IHI in ppTMS-EMG measures; however, the first phase (10 ms) could not be compared with spTMS-EEG due to the large TMS artefacts. In our study, IHI peaked at an ISI of 30 ms, which corresponded with a large positive peak around 30 ms in TEPs over the right contralateral M1. This is in agreement with the results of source estimations suggesting that following suprathreshold stimulations over M1, TEPs between 20 and 40 ms mainly originate from the contralateral M1 [74, 75]. Despite some weak evidence for relationships, interhemispheric EEG and EMG measures of cortical inhibition did not strongly correlate in amplitude or shape, in the time or frequency domain. In our previous study, we found that TEPs over contralateral electrodes are highly correlated with PEPs in the time domain [15]. In the current study, we extend this finding to the frequency domain, demonstrating high correlations between TEPs and PEPs in both low and high-frequency bands over contralateral electrodes. Therefore, it is likely that contralateral TEPs and TMS-evoked oscillations are highly sensitive to sensory co-activation, which could mask any signal propagating from the stimulated motor cortex. In sum, the findings did not provide strong evidence relating the EMG and EEG measures of TMS-evoked interhemispheric inhibition, possibly due to the sensitivity of contralateral TEPs to sensory activity resulting from the TMS pulse.

## 5. Conclusions

In contrast to our hypotheses, we did not find evidence for a relationship between the conditioned MEPs following ppTMS and TEPs following spTMS in the time domain. Instead, we found a similar time course of fluctuations in the magnitude of high-frequency oscillations in TEPs and the amplitude of conditioned MEPs, regardless of TMS intensity or waveform, providing preliminary evidence that the two measures reflect excitability from a similar cortical population. We also found that both high- and low-frequency oscillations in TEPs were correlated with TMS-evoked sensory activity, particularly over contralateral electrodes, and the amplitude of both TEPs and conditioned MEPs (from ISIs ~30 ms) were influenced by the size of TMS-evoked muscle contractions following a conditioning pulse. These findings highlight that both TEPs and conditioned MEPs are sensitive to peripheral factors other than the excitability of the stimulated cortical region or network. As such, care must be taken when interpreting or comparing outcomes from TMS-EEG and ppTMS-EMG studies.

## Acknowledgments

This research was supported by the National Health and Medical Research Council of Australia (1072057, NR; 1104580, AF, NR) and the Australian Research Council (180100741, NR).

