## Supplementary Materials for "The correspondence between EMG and EEG measures of changes in cortical excitability following transcranial magnetic stimulation"

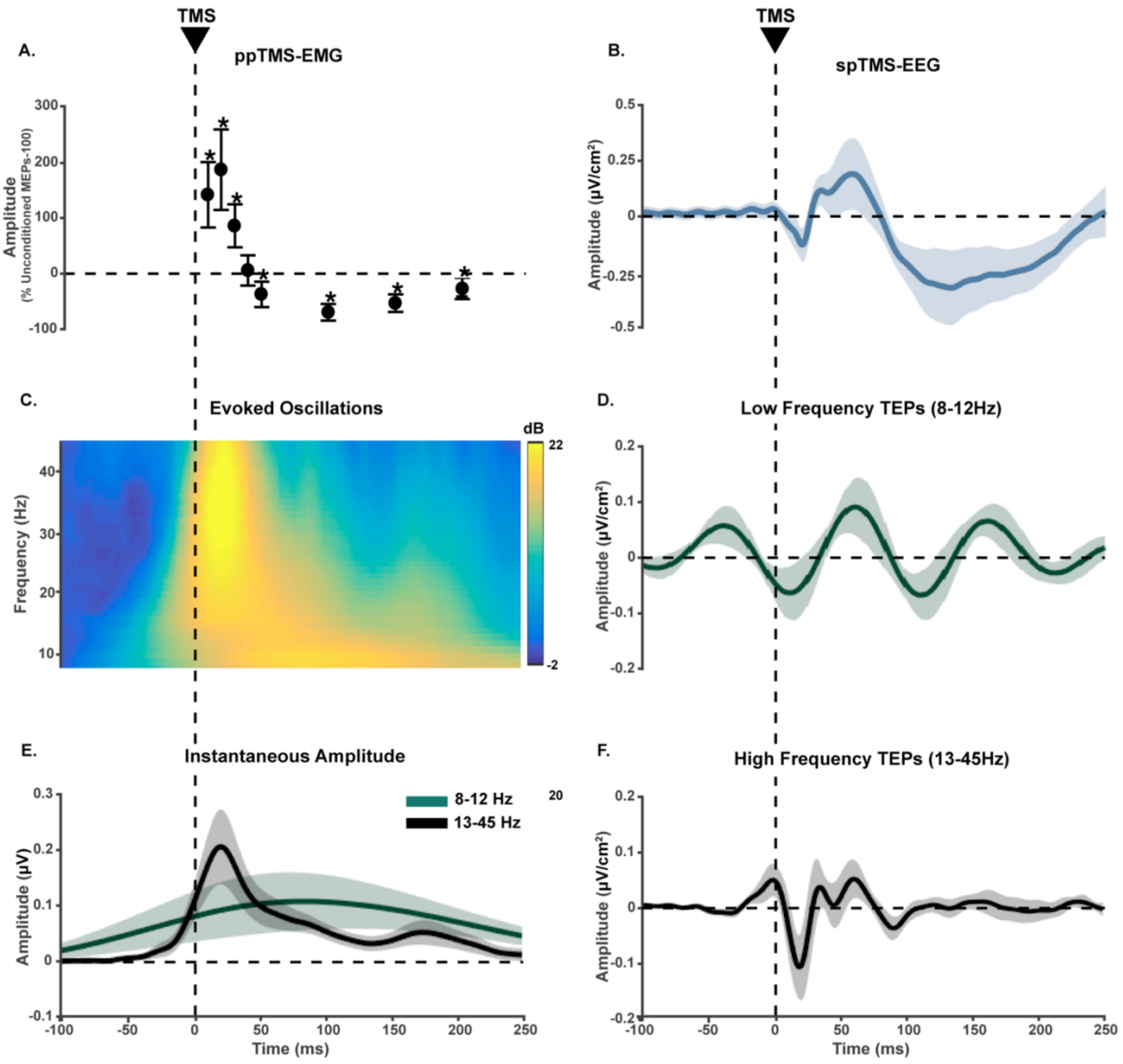


**Figure S1**: **EMG and EEG measures of local cortical responses to suprathreshold Monophasic TMS.** A) ppTMS-EMG recordings. Both conditioning stimulus and test stimulus were administered to left M1 and MEPs were recorded from right FDI. The solid circles represent the mean amplitude of conditioned MEPs expressed as a percentage of the mean unconditioned MEP amplitude (across individuals) from TS alone. The results are subtracted by 100 to show inhibition as a negative value. The error bars indicate 95% of confidence intervals assuming a Gaussian distribution. *indicate $p < 0.05$ comparing conditioned and unconditioned MEPs. B) spTMS-EEG recordings in the time domain. TMS is applied over left M1 with the same intensity as the conditioning stimulus in ppTMS. TEPs show the average of recordings at C3, FC1, CP1, FC5, CP5, C1, FC3, CP3, C5. The shaded areas show the 95% of confidence intervals. C) Distribution of the evoked oscillatory amplitude of different frequency bins recorded at left M1 across time, obtained from Morlet wavelet decomposition. D-F) TEPs filtered into the frequency bands with maximum power (observed in C). E) Instantaneous amplitude of the signals filtered into low and high frequencies. The line graphs show the group average of Hilbert amplitude changes recorded at left M1 and the shaded areas represent the 95% of confidence intervals.

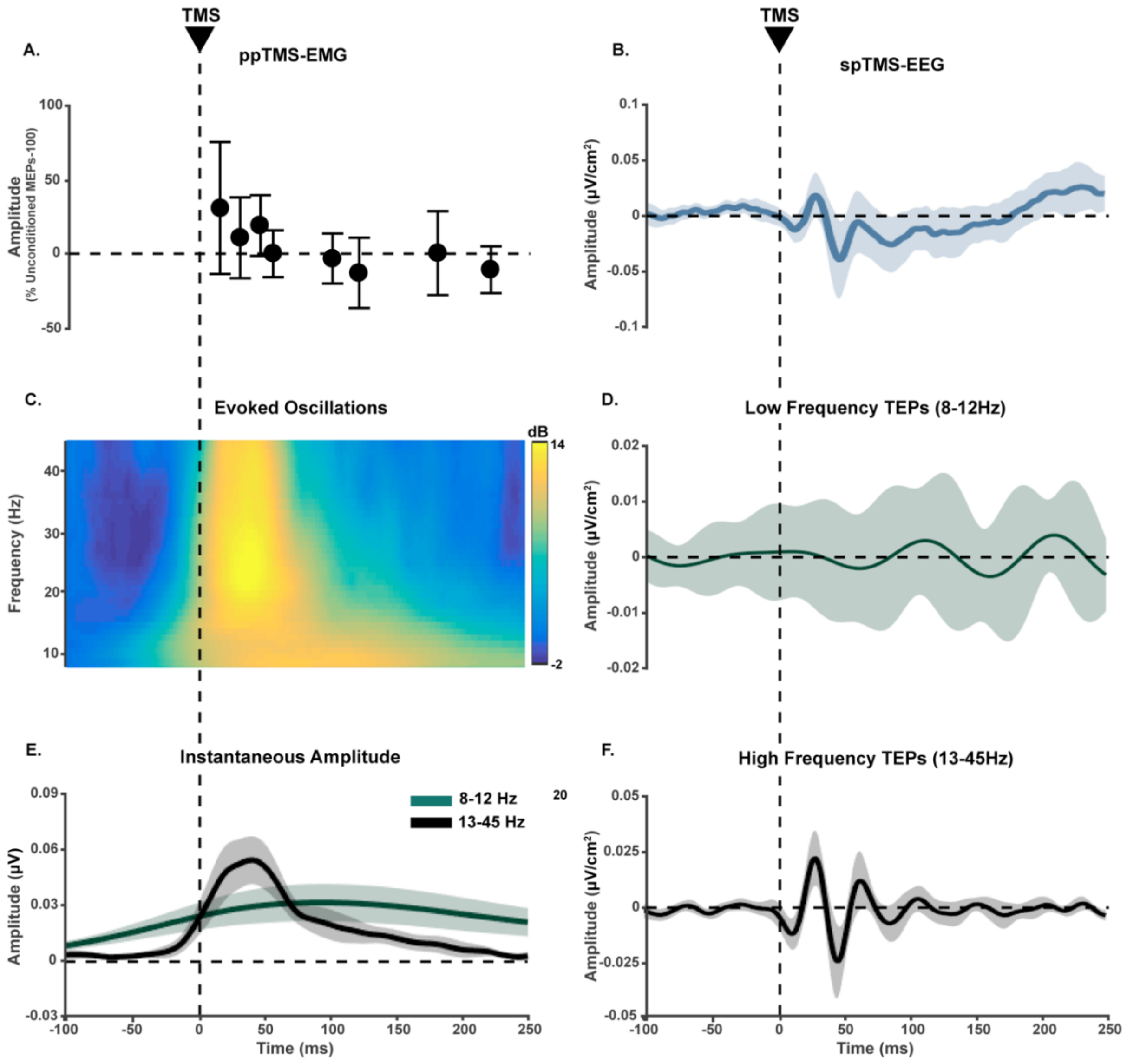


**Figure S2**: **EMG and EEG measures of local cortical responses to subthreshold biphasic TMS.** A) ppTMS-EMG recordings. Both conditioning stimulus and test stimulus were administered to left M1 and MEPs were recorded from right FDI. The solid circles represent the mean amplitude of conditioned MEPs expressed as a percentage of the mean unconditioned MEP amplitude (across individuals) from TS alone. The results are subtracted by 100 to show inhibition as a negative value. The error bars indicate 95% of confidence intervals assuming a Gaussian distribution. *indicate $p < 0.05$ comparing conditioned and unconditioned MEPs. B) spTMS-EEG recordings in the time domain. TMS is applied over left M1 with the same intensity as the conditioning stimulus in ppTMS. TEPs show the average of recordings at C3, FC1, CP1, FC5, CP5, C1, FC3, CP3, C5. The shaded areas show the 95% of confidence intervals. C) Distribution of the evoked oscillatory powers of different frequency bins recorded at left M1 across time, obtained from Morlet wavelet decomposition. D-F) TEPs filtered into the frequency bands with maximum power (observed in C). E) Instantaneous amplitude of the signals filtered into low and high frequencies. The line graphs show the group average of Hilbert amplitude changes recorded at left M1 and the shaded areas represent the 95% of confidence intervals.

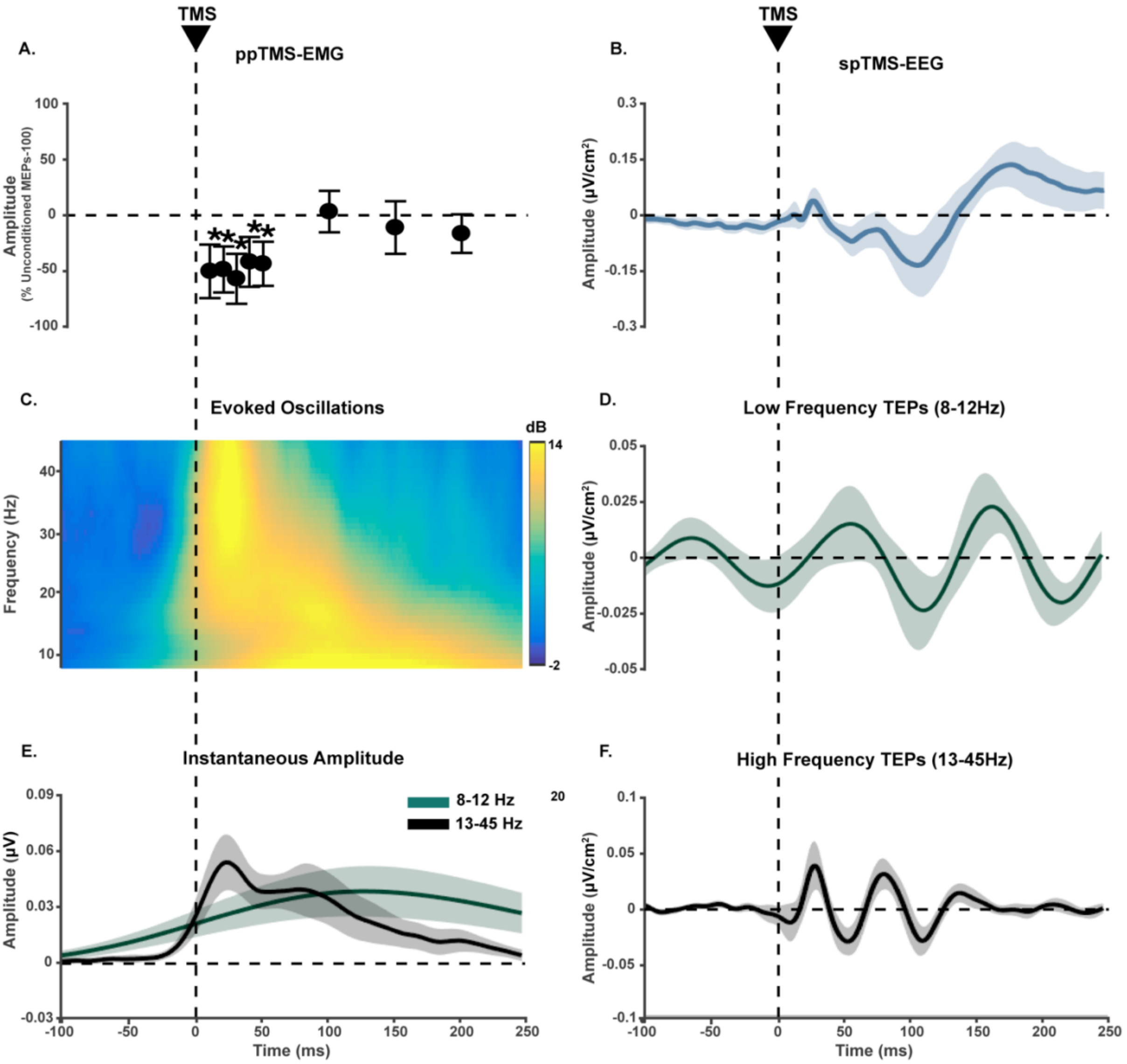


**Figure S3**: **EMG and EEG measures of interhemispheric cortical responses to suprathreshold monophasic TMS.** A) ppTMS-EMG recordings. Conditioning and test stimulus were administered to left and right M1s, respectively, and MEPs were recorded from left FDI. The solid circles represent the mean amplitude of conditioned MEPs expressed as a percentage of the mean unconditioned MEP amplitude (across individuals) from TS alone. The results are subtracted by 100 to show inhibition as a negative value. The error bars indicate 95% of confidence intervals assuming a Gaussian distribution. *indicate $p < 0.05$ comparing conditioned and unconditioned MEPs. B) spTMS-EEG recordings in the time domain. TMS is applied over left M1 with the same intensity as the conditioning stimulus in ppTMS. TEPs show the average of recordings at C4, FC2, CP2, FC6, CP6, C2, FC4, CP4, and C6. The shaded areas show the 95% of confidence intervals. C) Distribution of the evoked oscillatory powers of different frequency bins recorded at right M1 across time, obtained from Morlet wavelet decomposition. D-F) TEPs filtered into the frequency bands with maximum power (observed in C). E) Instantaneous amplitude of the signals filtered into low and high frequencies. The line graphs show the group average of Hilbert amplitude changes recorded at right M1 and the shaded areas represent the 95% of confidence intervals.

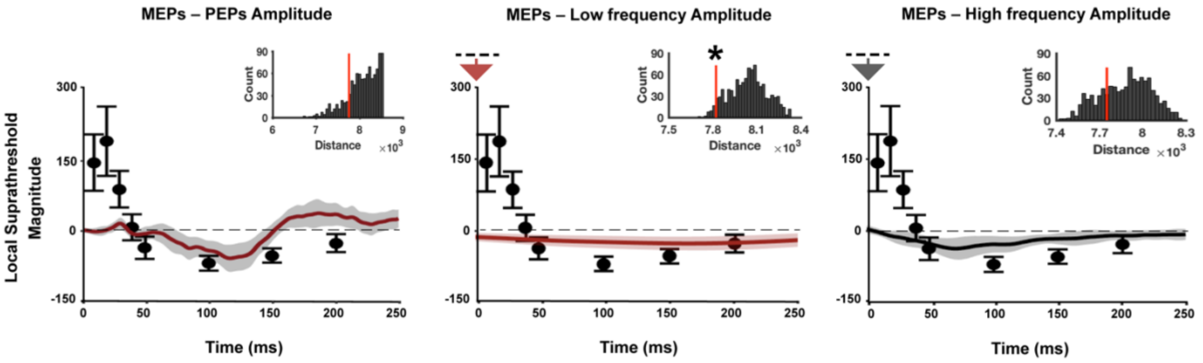


**Figure S4: Shape similarity between PEPs and EMG measures of cortical excitability in local responses to suprathreshold and monophasic TMS.** The line graphs represent group-averaged EEG recordings at left M1 (C3, FC1, CP1, FC5, CP5, C1, FC3, CP3 and C5) to spTMS over left shoulder. The shaded areas show the 95% of confidence intervals. Measures of oscillatory amplitude are baseline-corrected (-600 to -100ms). The solid circles represent the group-averaged changes in MEPs (conditioned relative to unconditioned) recorded from right FDI in response to ppTMS over left M1. The error bars indicate 95% of confidence intervals assuming a Gaussian distribution. The embedded bar plots depict the result of shape-similarity tests between the two signals. * shows significant similarity in shape between the two signals ($p<0.05$). The downward arrow on the top of Y axis indicates that EEG measures are rescaled to negative values.
